# Two avian *Plasmodium* species trigger different transcriptional responses on their vector *Culex pipiens*

**DOI:** 10.1101/2023.01.24.525339

**Authors:** M Garrigós, G Ylla, J Martínez-de la Puente, J Figuerola, MJ Ruiz-López

## Abstract

Malaria is a mosquito-borne disease caused by protozoans of the genus *Plasmodium* that affects both humans and wildlife. The fitness consequences of infections by avian malaria are well known in birds, however, little information exists on its impact on mosquitoes. Here we study how *Culex pipiens* mosquitoes transcriptionally respond to infection by two different *Plasmodium* species, *P. relictum* and *P. cathemerium*, differing in their virulence (mortality rate) and transmissibility (parasite presence in exposed mosquitoes’ saliva). We study the mosquito response to the infection at three critical stages of parasite development: formation of ookinetes at 24 hours post-infection (hpi), the release of sporozoites into the hemocoel at 10 days post-infection (dpi), and storage of sporozoites in the salivary glands at 21dpi. For each time point, we characterized the gene expression of mosquitoes infected with each *P. relictum* and *P. cathemerium* and mosquitoes fed on an uninfected bird and, subsequently, compared their transcriptomic responses. Differential gene expression analysis showed most of the transcriptomic changes occurred during the early infection stage (24 hpi), especially when comparing *P. relictum* and *P. cathemerium* infected mosquitoes. Differentially expressed genes in mosquitoes infected with each species were related mainly to the immune response, trypsin, and other serine-proteases metabolism. We conclude that these differences in response may partly play a role in the differential virulence and transmissibility previously observed in *P. relictum* and *P. cathemerium* in *Cx. pipiens*.

## 1. Introduction

Vector-borne diseases are a major challenge for both human and animal health, representing more than 17% of emerging infectious diseases (WHO 2021). Mosquitoes are vectors of relevant pathogens including viruses that cause yellow fever, dengue, or West Nile fever and parasites such as nematode worms which cause lymphatic filariasis, and haemosporidians, like *Plasmodium*, which cause malaria (Lehane 2010). Malaria is one of the most important vector-borne diseases for humans, and only in 2020, it caused around 602,000 deaths (WHO 2021). Malaria parasite species affect humans and related primates as well as other mammals, reptiles, or birds, driving some populations to extinction (van Riper et al. 1986).

Avian malaria is a worldwide distributed mosquito-borne disease caused by *Plasmodium* parasites which use birds as obligate hosts (Valkiūnas 2005). Avian *Plasmodium* is transmitted by mosquitoes, mainly of the genus *Culex,* which are the definitive hosts and where *Plasmodium* reproduces sexually (Valkiūnas 2005). In birds, as a part of their life cycle, *Plasmodium* merozoites invade the host erythrocytes and can differentiate into mature gametocytes (Sinden 1983). When a female mosquito feeds on a *Plasmodium*-infected bird, gametocytes are released from the erythrocytes, and male and female gametes unite resulting first in the zygote, and then in the motile ookinete (Sinden 2002). About 24 hours post-infection (hpi) the ookinete crosses the midgut epithelium and remains located between the epithelial surface and the basal lamina, where it transforms into a sessile oocyst. After about 10 days post-infection (dpi), mature oocysts liberate thousands of sporozoites into the hemocoel. Sporozoites that survive the immune system of the mosquito eventually invade the salivary glands (Sinden 1983; Sinden 2002; Abraham and Jacobs-Lorena 2004) from where they can be then transmitted to a new avian host upon a mosquito bite.

For an appropriate development and transmission of the parasites, mosquitoes susceptible to the infection need to survive it, allowing parasites to complete their life cycle. Thus, the nature of the interaction between pathogens and mosquitoes determines the ability of a vector to acquire, maintain and transmit parasites to a new host, i.e. the vector competence (Beerntsen et al. 2000; Bonizzoni et al. 2013). Therefore, the vector competence is conditioned on the mosquito response against the pathogen (Higgs and Beaty 2005), which includes a number of defense mechanisms. After a blood meal, mosquitoes synthesize and release serine proteases that constitute a chemical barrier against pathogens (Vizioli et al. 2001; Molina-Cruz et al. 2005; Muller et al. 1995) and form a chitin-containing peritrophic matrix around the blood. This matrix constitutes a physical barrier for harmful food particles, digestive enzymes, and pathogens (Lehane 1997). Despite the existence of these and other barriers, some pathogens manage to reach the mosquito midgut, hemocoel, or internal organs and activate the immune response, which may be categorized into cellular and humoral immune responses (Hillyer 2016). The cellular response includes mechanisms such as phagocytosis, cellular encapsulation, autophagy, melanization, and induction of apoptosis, while the humoral response consists of the activation of signaling pathways that eventually result in the synthesis of factors with antimicrobial activity (Michel and Kafatos 2005; Hillyer 2016). In insects, one of the main immune signaling pathways is the Toll pathway, which is activated by the Spätzle cytokine and results in the activation of the transcription of antimicrobial peptides (AMPs) and other immune effectors that may combat pathogens (Kumar et al. 2018). In addition, two other pathways have been shown to be involved in the response to *Plasmodium*, including avian malaria parasites, the immune deficiency (Imd) and the Janus Kinase signal transducer of activation (JAK- STAT) (Clayton et al. 2014, García-Longoria et al. 2022).

The infection by *Plasmodium* and the mosquito response against infection ultimately result in a cost to vectors (Ahmed et al. 2002), which might depend, among other factors, on which *Plasmodium* species infects the mosquito. For example, Gutiérrez-López et al. (2020), found a higher survival rate and transmissibility (measured as the presence of parasite DNA in the saliva of mosquitoes) in *Culex pipiens* infected with *Plasmodium cathemerium* compared to those infected with *Plasmodium relictum*. However, the genetic mechanisms that underlay these phenotypic differences are unknown. Part of the heterogeneity in parasite virulence and therefore in the fitness consequences in their hosts (Gutiérrez-López et al. 2020) is due to the remarkable cellular plasticity and transcriptional variation of *Plasmodium* (García-Longoria et al. 2020). Avian *Plasmodium* is an extremely diverse clade with at least 55 species (Valkiūnas and Iezhova 2018) divided into more than 1,446 unique genetic lineages (Bensch et al. 2009). This extensive inter and intra-specific genomic variation will translate into different phenotypic characters, including differences of virulence, which will also determine how the mosquitoes respond to an infection.

Although studies on avian malaria that use transcriptomic approaches are increasing in the last few years, they have mainly focused on the avian host, addressing either the bird response to infection (e.g. Videvall et al. 2015) or the gene expression of *Plasmodium* infecting birds (e.g. Videvall et al. 2017; García-Longoria et al. 2020; Videvall et al. 2021). In contrast, to the best of our knowledge, only three studies have analysed the gene expression of mosquitoes infected by avian *Plasmodium* (Zou et al. 2011; Ferreira et al. 2022; García-Longoria et al. 2022). However, none of them focus on *Cx. pipiens*, which is the main vector of avian *Plasmodium* in Europe. Another significant limitation is that all of these studies focus on *P. relictum* and consequently do not take into consideration the effect of the differences in virulence and transmissibility between *Plasmodium* lineages.

Here, we analyse the gene expression of *Cx. pipiens* infected with two widely distributed species of avian *Plasmodium* with different characteristics namely, *P. relictum* (lineage SGS1) and *P. cathemerium* (lineage PADOM02). To that end, we obtained RNA-seq data of mosquitoes infected with the two *Plasmodium* species at three time points corresponding with key stages of parasite development in the vector. These stages are (1) during the formation of ookinetes, (2) during the release of sporozoites into the hemocoel, and (3) after sporozoites invade and are stored in the salivary glands. This study uses a natural avian malaria system that will help to address the current knowledge gaps on molecular mechanisms occurring during *Plasmodium* infections in mosquitoes. In addition, these results will allow us to further understand the differences in *Cx. pipiens* response against different *Plasmodium* species and how this might affect vector competence.

## 2. Material and Methods

### Sampling and experimental conditions

We captured juvenile house sparrows (*Passer domesticus*) with mist nets in September 2020 at Granja Escuela de Trigueros (Huelva province, Spain). We ringed, weighted and measured the individuals before bringing them into captivity at the animal facilities of the Doñana Biological Station, following the ethical guidelines (article 34 RD 53/2013).

In the field, we took blood samples from each bird jugular vein using a sterile syringe, taking both blood samples and blood smears. Blood samples were used to visually and molecularly identify the blood parasite infections and the parasite lineage identity. To do that, we extracted genomic DNA from blood samples using a Lithium Chloride protocol (Gemmell and Akiyama 1996) and detected parasite infections following Hellgren et al. (2004). We sequenced the amplified products for positive samples on both strands using Capillary Electrophoresis Sequencing by Macrogen (Madrid, Spain). We analysed the sequences using Geneious v. 2020.0.3 (Kearse et al. 2012) and identified lineages in MalAvi (Bensch et al. 2009). After molecular analyses, we chose three birds for further analyses, namely: A bird that was not infected by *Plasmodium*, *Haemoproteus* nor *Leucocytozoon* (control), a bird infected with *P. relictum* lineage SGS1, and a bird infected with *P. cathemerium* lineage PADOM02. To visually assess the intensity of parasitemia we analyzed the blood smears of the selected birds infected by *P. cathemerium* and *P. relictum* using a light microscope Olympus CX33. The intensity of parasitemia was less than 1 parasite cell/15.000 erytrocites in both cases.

We collected mosquito larvae on October 2020 in Aljaraque (Huelva province) and reared them following Gutiérrez-López et al. (2020). We maintained larvae in dechlorinated water and fed *ad libitum* with Hobby-Mikrozell 20 ml/22 g (Dohse Aquaristik GmbH & Co.101 KG, D-53501, Gelsdorf, Germany) and Hobby-Liquizell 50 ml (Dohse Aquaristik GmbH & Co.101 KG, D-53501, Gelsdorf, Germany). After emergence, we identified adults to the species level and sexed them following Gunay et al. (2018). We kept adult *Cx. pipiens* females in separate cages of 50 individuals maximum and fed them with a 10% sugar solution. We maintained both larvae and adult mosquitoes under controlled conditions (26°C ± 1, 55-60% relative humidity (RH) and 12:12 light:dark photoperiod cycle).

We divided the 11-day-old adults (± 1 day) into 3 groups mixing individuals originating from the different cages and allowed them to feed overnight on a *P. relictum*-infected bird, a *P. cathemerium*-infected bird and an uninfected control bird. Only one individual of each category was used in this experiment and they were exposed to mosquitoes only one time. In the morning after exposure, we separated the fed females of each group into three different cages and maintained them under the conditions described above.

For transcriptome analyses, we processed mosquitoes at three time points after exposure (24 hpi, 10 dpi and 21 dpi). At each time point, we created pools of 5 mosquitoes of each infection status capturing the mosquitoes alive and immediately transferring them to dry ice. To ensure that 24h had passed since feeding, the degree of blood digestion was assessed according to the Sella scale, following Detinova et al. 1962., and selecting mosquitos classified as Sella stage IV. We preserved the mosquitoes at -80C until RNA extractions were carried out. We collected a total of 36 samples including controls (4 pools x 3 time-points x 3 conditions).

### RNA extraction, library preparation, and sequencing

We extracted RNA and DNA from pools of 5 mosquitoes using TRIzol® (Invitrogen, Carlsbad, CA, USA) followed by column purification using RNeasy mini kit® (QIAGEN, Hilden, Germany) following Ferreira et al. (2022). We tested the remaining DNA for the presence of *Plasmodium* following Hellgreen et al. (2004) confirming that parasite DNA was present in all positive samples and not present in negative samples. We prepared RNAseq libraries at the Polo d’Innovazione di Genomica, Genetica e Biologia, Siena (Italy) using and Illumina library. Then, we quantified samples using Qubit® 4.0 Fluorometer and checked RNA integrity using the Fragment Analyzer to measure RNA Quality Number (RQN) specific for mosquitoes. Finally, we prepared the libraries following the QIAseqTM Stranded mRNA Selected Kit Handbook for Illumina Paired-End Indexed Sequencing. Indexed DNA libraries were sequenced in an Illumina NextSeq550 Flowcell, using the Illumina chemistry V2.5, 2x75bp run.

### Data analysis

To check the quality of the reads we used FastQC (ver. 0.11.9; Andrews et al. 2010) and MultiQC (Ewels et al. 2016). Then, we filtered low quality and under 36 bp reads using Trimmomatic (Bolger et al. 2014). Because the reference genome and annotations of *Cx. pipiens* are not published yet, we used the reference genome and annotations of phylogenetically closest species that were available in Ensembl, *Cx. quinquefasciatus* (https://metazoa.ensembl.org/Culex_quinquefasciatus/Info/Index). We used STAR (Dobin et al. 2012) to map the short reads to the reference genome of *Cx. quinquefasciatus* and RSEM (Li and Dewey 2011) to quantify gene abundances.

Following steps were carried out in R (R Core Team 2021) using Bioconductor packages (Huber et al. 2015). We carried out a Variance Stabilizing Transformation (VST) of the counts to represent the samples on a PCA plot. Then, we used the DESeq2 package (Love et al. 2014) to perform the differential gene expression analysis comparing: i) *P. cathemerium* infected mosquitoes vs controls, ii) *P. relictum* infected mosquitoes vs controls, and iii) *P. relictum* infected mosquitoes vs *P. cathemerium* infected mosquitoes. We kept those genes with an adjusted p-value < 0.01 and sorted them by the log2 fold changeestimations, considering genes with a LFC>1 or LFC<-1, excepting one gene specified above. We finished the differential gene expression analysis visualizing differentially expressed genes by MA plots and an UpSet plot.

Finally, we performed a Gene Ontology (GO) enrichment analysis for the up- and down-regulated genes using the topGO package (Alexa and Rahnenfuhrer 2010) including the “biological processes” and “molecular functions” categories from VectorBase (Giraldo-Calderón et al. 2015). For the Enrichment analysis we used classical algorithm and Fisher’s exact test and considered enriched the GO terms with p- value < 0,01. We used the ggplot2 (Wickham 2016) package to visualize the enriched GO terms as described by Bonnot et al. (2019).

## 3. Results

### Transcriptomics data description

We obtained 36 mRNA-seq libraries: four replicates for three infection status at three time points. The mean number of reads per library was 33,177,082, ranging from 15,406,828 to 45,332,626 reads for raw samples. Raw data has been made publicly available through to the European Nucleotide Archive ENA database (https://www.ebi.ac.uk/ena/browser/home) under project accession number PRJEB1609, Study ERP125411. The percentage of reads kept after trimming ranged from 80.56% to 99.52%, (12,411,723 to 44,887,231 reads see Table S1). The MultiQC report showed a mean quality score above q30 in all base calls across the read. On average, 80.87% of reads mapped to the genome of *Cx. quinquefasciatus*. For downstream analyses, we removed two samples taken at 24 hpi that had barcoding errors due to lab processing.

### Sample clustering reveals early transcriptomic response to infection

The principal component analysis (PCA) results show that most of the transcriptome variation is contained in the PC1 (85% var.) driven by the time post-bloodmeal, under both *Plasmodium* species infections. At 24 hpi, there were clear differences (PC2, 3% var.) between control samples, *P. cathemerium*, and *P. relictum* infected mosquitoes. By contrast, there were no clear transcriptomic differences at later stages, such as between 10 dpi and at 21 dpi, nor between infection statuses (Figure 1).

**Figure 1.**
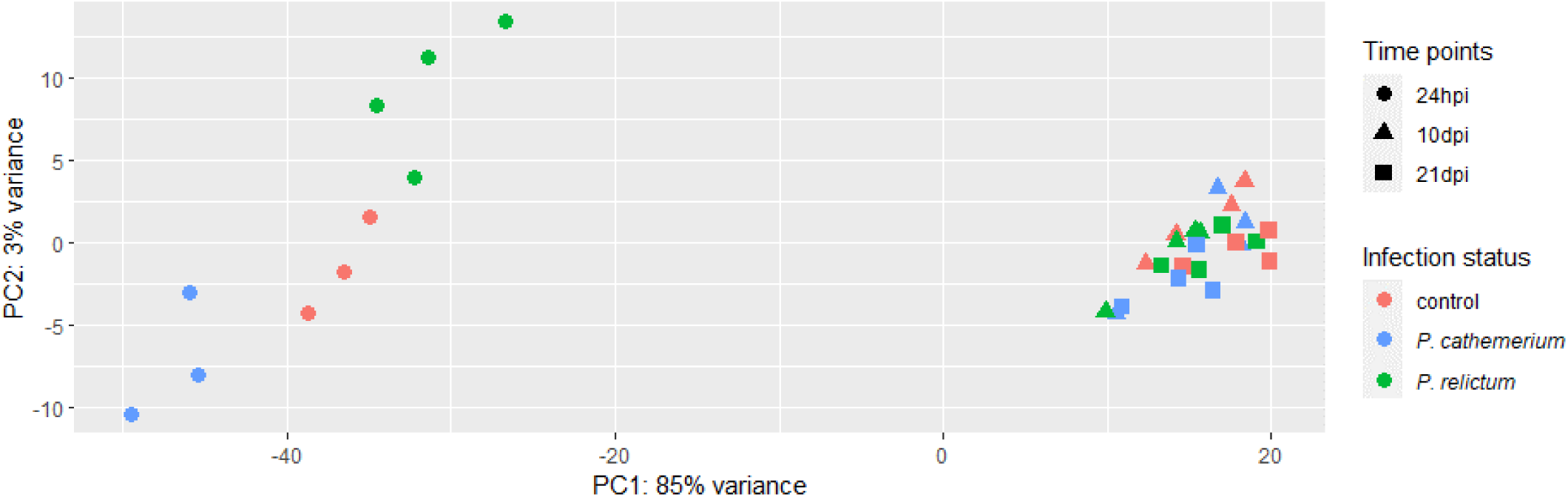
Principal component analysis (PCA) of transcriptome variation *in Cx. pipiens* at three time points after feeding on either an uninfected bird (control), a *P. cathemerium*-infected bird and a *P. relictum*-infected bird. Time points analysed were 24 hours post-infection (hpi), 10 days post-infection (dpi) and 21 dpi. The x-axis shows the first principal component score, which captures 84% of variation and the y-axis shows the second principal component score, which captures 3% of variation.

### Differential gene expression analysis and enrichment analysis unveil the clues of the mosquitoe transcriptomic response to Plasmodium infections

Overall, 2,051 genes were differentially expressed in *Cx. pipiens,* from which 590 showed a log2 fold change (LFC) higher than 1 or lower than -1 (see Table 1 for differentially expressed genes distribution among comparisons). Most of the transcriptomic differences were found at an early infection stage (24 hpi), especially when comparing *P. relictum*-infected mosquitoes vs *P. cathemerium-*infected mosquitoes, and *P. relictum-*infected mosquitoes vs controls. No differences were found between mosquitoes infected by *P. cathemerium* and those infected by *P. relictum* at 10 dpi and between mosquitoes infected with *P. relictum* and controls at 21 dpi. At 24 hpi, the comparisons *P. relictum*-infected mosquitoes vs controls and *P. relictum* vs *P. cathemerium*-infected mosquitoes shared 460 differentially expressed genes (Figure 2; see Supplementary Material for MA plots).

**Figure 2.**
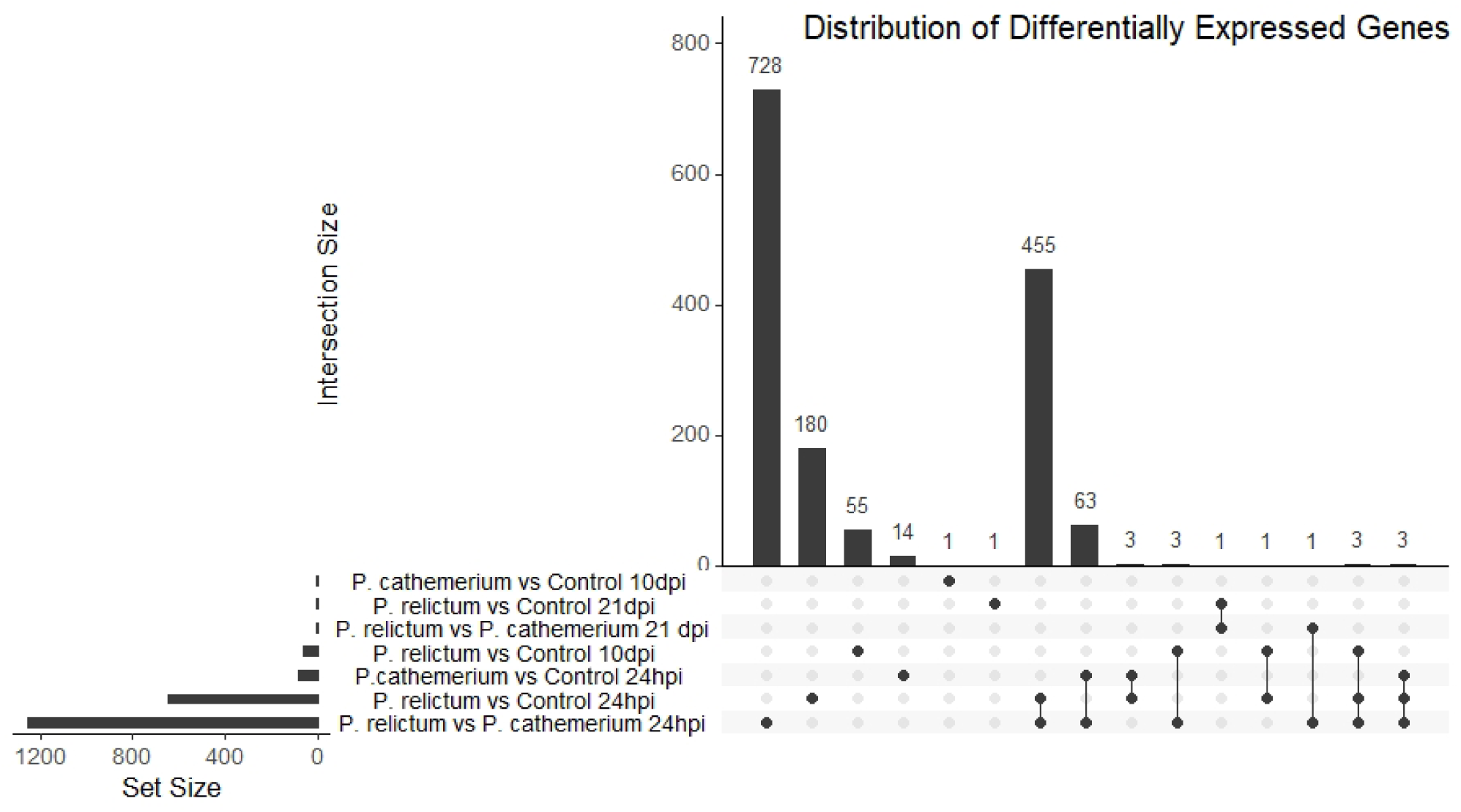
UpSet plot showing overlap size of sets of differentially expressed genes for i) *P. cathemerium* infected mosquitoes vs controls, ii) *P. relictum infected* mosquitoes vs controls, and iii) *P. relictum* infected mosquitoes vs *P. cathemerium* infected mosquitoes at three time points (24 hpi, 10 dpi and 21 dpi). The top vertical bar plot shows the number of genes (y-axis) contained in each intersection (x-axis). The horizontal bar plot at the bottom shows the number of differentially expressed genes for each comparison.

**Table 1.**
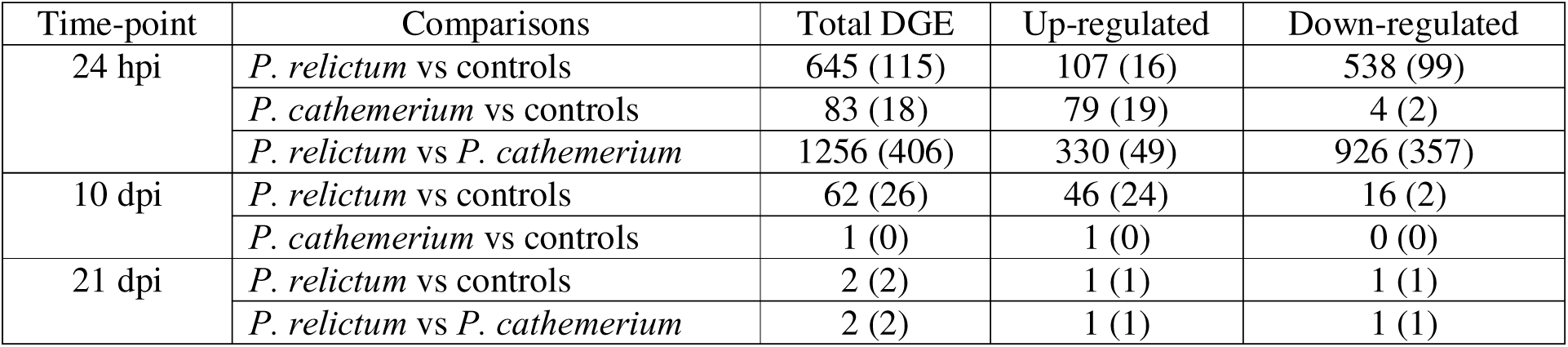
Number of differentially expressed genes (DEG) with a p<0.01 for all comparisons at 24 hours post-infection (24 hpi), 10 days post-infection (10 dpi), and 21 days post-infection (21 dpi). In brackets, DEG filtered by log2 fold change (LFC) > 1 for up-regulated genes and LFC < -1 for down-regulated genes. No DEG were found for the comparisons *P. relictum* vs *P. cathemerium* infected mosquitoes at 10 dpi and *P. cathemerium* infected mosquitoes vs controls at 21 dpi.

#### a) *Cx. pipiens* gene expression response to *P. relictum* infection

At 24 hpi, up-regulated genes in *P. relictum*-infected *Cx. pipiens* compared to controls included a cecropin gene (CPIJ005108), which is part of the Toll pathway in the immune response, and a mitochondrial NADH-ubiquinone oxidoreductase gene (CPIJ009076). Another gene involved in the Toll pathaway, *spätzle 1B* (CPIJ006792), was also up-regulated in *P.relictum*-infected mosquitoes, although with a LFC = 0.3984. Within the down-regulated genes there were four multicopper oxidase genes (CPIJ016802, CPIJ010466, CPIJ012244, and CPIJ010465), four chymotrypsins (CPIJ003915, CPIJ018205, CPIJ007838, and CPIJ006568) and two serine proteases genes (CPIJ004984 and CPIJ002112) (Figure S1A).

At 10 dpi, when compared to controls, mosquitoes infected by *P. relictum* two up-regulated genes were ribosomal genes (CPIJ040837 and CPIJ009519). At 21 dpi, the aromatic amino acid decarboxylase gene (CPIJ010562) was down-regulated (Figure S1B).

At 24 hpi, the GO categories within biological processes related to cellular mechanisms and mitochondrial chain complex assemblies presented the greatest number of up-regulated genes. Within down-regulated genes, the enriched biological processes included molecules metabolism, transport and localization (Figure 3). The molecular functions enriched for up-regulated genes were cation binding, metal ion binding and ion binding. Protein, ATP, and nucleic acids binding molecular functions (including purine ribonucleoside triphosphate, purine nucleotide, purine ribonucleotide, adenyl nucleotide and adenyl ribonucleotide, among others) were enriched for down-regulated genes (Figure 4).

**Figure 3.**
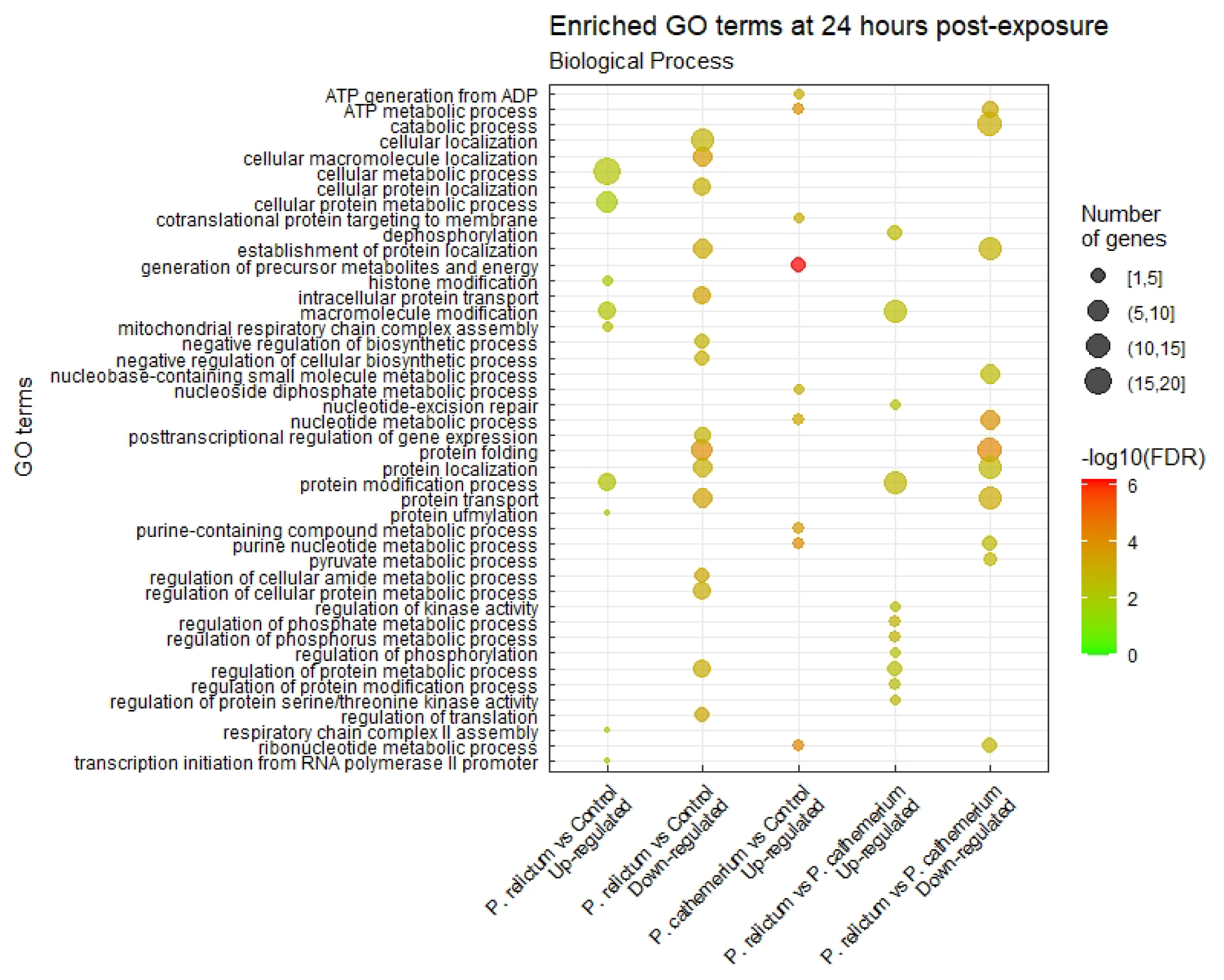
Dot plot of significantly (adjusted p-value <0.01) enriched GO biological processes for differentially expressed genes (up and down-regulated) at 24 hpi for i) *P. cathemerium* infected mosquitoes vs controls, ii) *P. relictum* infected mosquitoes vs controls, and iii) *P. relictum* infected mosquitoes vs *P. cathemerium* infected mosquitoes. We did not find enriched GO terms for down-regulated genes for *P. cathemerium* infected mosquitoes vs controls. Larger dots correspond to a higher number of significant genes, and the color gradient goes from green for the least significant terms to red for the most significant terms, measured as False Discovery Rate (FDR).

**Figure 4.**
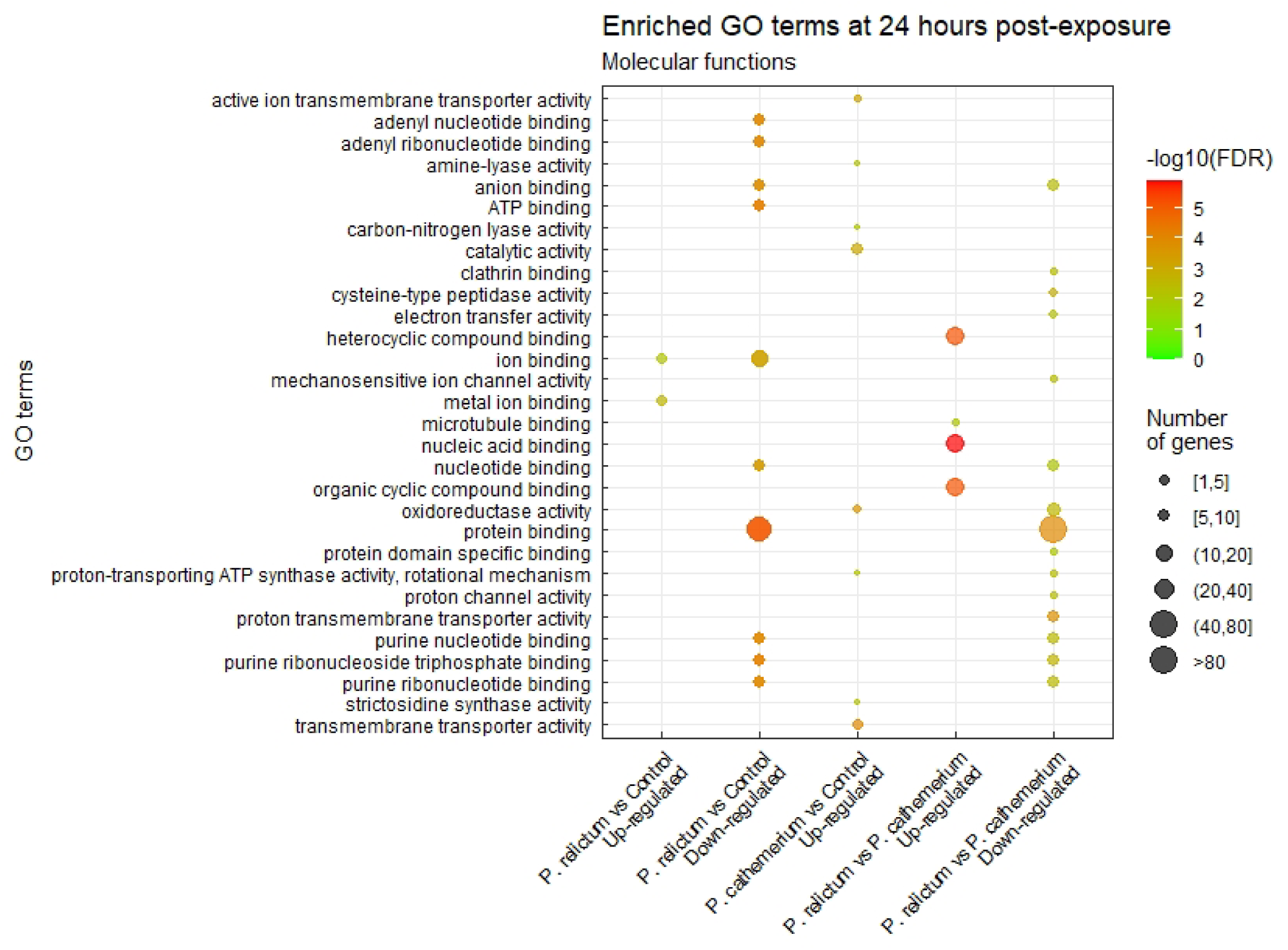
Dot plot of significantly (adjusted p-value <0.01) enriched GO molecular functions for differentially expressed genes (up and down-regulated) at 24 hpi for i) *P. cathemerium* infected mosquitoes vs controls, ii) *P. relictum* infected mosquitoes vs controls, and iii) *P. relictum* infected mosquitoes vs *P. cathemerium* infected mosquitoes. We did not find enriched GO terms for down-regulated genes for *P. cathemerium* infected mosquitoes vs controls.

At 10 dpi enriched biological processes were related to amino acid biosynthesis for up-regulated genes and to protein geranylgeranylation for the down-regulated genes (Figure S2). Molecular functions included actin monomer binding and GTPase activity for up-regulated genes (Figure S3).

#### b) *Cx. pipiens* gene expression during response to *P. cathemerium* infection

In *P. cathemerium*-infected mosquitoes compared to controls at 24 hpi (Figure S4), we found up-regulated genes related to digestive enzymes like *trypsin 2 precursor* (CPIJ005273), two protein G12 precursors (CPIJ012846 and CPIJ012848), and genes related to the mitochondrial electron chain including genes for an ADP-ATP carrier protein (CPIJ005941), a mitochondrial phosphate carrier protein (CPIJ013141), and the cytochrome P450 4C51 (CPIJ018944). The *pom1* gene (CPIJ019938) was down-regulated. At both 10 dpi and 21dpi no genes had differential gene expression with LFC > 1.

At 24 hpi, GO terms associated with biological processes were mostly related to the generation of precursor metabolites and energy, ATP generation, and nucleic acids metabolic processes, including nucleoside phosphate, ribose phosphate, nucleotide, ribonucleotide, purine nucleotide and purine ribonucleotide metabolic processes (Figure 3). GO terms associated with molecular functions included ion and transmembrane transporter, oxidoreductase and proton-transporting ATP synthase activities (Figure 4).

#### c) *Cx. pipiens* respond differently to *P. relictum* and *P. cathemerium* infections

At 24 hpi, the comparison *P. relictum* vs *P. cathemerium-* infected mosquitoes showed the larger number of differentially expressed genes (Table 1; Figure S5). The up-regulated genes included four zinc finger proteins genes (CPIJ015530, CPIJ012592, CPIJ008782, and CPIJ009347). On the other hand, mosquitoes infected by *P. relictum* compared to those infected by *P. cathemerium* showed down-regulated genes of five protein G12 precursors (CPIJ005176, CPIJ012848, CPIJ012844, CPIJ012845, and CPIJ012846), four multicopper oxidases (CPIJ016802, CPIJ010466, CPIJ010465, and CPIJ000864), two chitin synthases (CPIJ014268 and CPIJ014269), a peritrophic membrane chitin binding protein (CPIJ007042), and a number of serine proteases including five chymotrypsin genes (CPIJ003915, CPIJ018205, CPIJ007838, CPIJ006542, and CPIJ006568) and three trypsin precursors (CPIJ005273, CPIJ006019, and CPIJ004660). From the down-regulated genes, three multicopper oxidases and three chymotrypsin precursors were also down-regulated in *P. relictum*-infected mosquitoes compared to controls and two G12 protein precursors and one trypsin precursor were also up-regulated in *P. cathemerium*-infected mosquitoes.

No differentially expressed genes were found at 10 dpi and only 2 genes at 21 dpi, a down-regulated testicular acid phosphatase precursor (CPIJ007697) and an up-regulated uncharacterized protein.

At 24 hpi, biological processes related to metabolic regulation were enriched for up-regulated genes, while down-regulated genes included nucleic acid nucleotide metabolic processes (Figure 3), some of them also enriched for up-regulated genes *in P. cathemerium*-infected mosquitoes compared to controls. Molecular functions like microtubule and nucleic acid binding were enriched in up-regulated genes in mosquitoes infected by *P. relictum* compared to those infected by *P. cathemerium*. Most down-regulated enriched molecular functions were related to protein and nucleotides binding (Figure 4), as observed in mosquitoes infected by *P. relictum* compared to controls.

## 4. Discussion

How mosquitoes respond to infection and the impact of such infection will ultimately influence the transmission of different *Plasmodium* species. For example, *P. relictum* and *P. cathemerium* infections in *Cx. pipiens* have a different impact on the mosquitoes’ fitness. In particular, *P. relictum* causes higher mortality than *P. cathemerium* infections in *Cx. pipiens*, and the transmissibility of *P. relictum* is lower than that of *P. cathemerium* (Gutiérrez-López et al. 2020). But little is known about the genetic underpinnings of mosquito response to *Plasmodium* infections. Here, we compare for the first time the transcriptional response of *Cx. pipiens* infected by these two avian *Plasmodium* species. Our results show that although responses of infected *Cx. pipiens* by these *Plasmodium* species share some common pathways, there are key differences in gene expression associated with the immune system and serine-protease synthesis. These information provides new insights into the molecular mechanism underlying *Culex pipiens* immune response against *Plasmodium*. Furthermore, by comparing two *Plasmodium* species we show that there is variation in the immune response of *Culex pipiens* to different avian malaria species. This information will in the long term help to understand the complex interaction between the mosquitos and different malaria parasites and potentially uncover the key to understanding differences on transmission and fitness consequences for the mosquitoes.

### Early and different response to the infection

When a mosquito feeds on a vertebrate host, a number of processes involving changes in gene expression are triggered (Dana et al. 2005). We found that the vast majority of the differentially expressed genes between infected and uninfected mosquitoes were found at 24 hpi. Between 18-24 h after blood feeding ookinetes form, invade the peritrophic matrix leaving behind the blood bolus and start the midgut epithelium cell invasion (Cirimotich et al. 2010; Valkiūnas et al. 2015; Baia-da-Silva et al. 2018). Ookinete formation and invasion of the midgut epithelium are considered critical steps that will determine the success of the infection. Parasite abundance drops drastically during this step due to lumenal and epithelial immune responses mounted by the mosquito (Cirimotich et al. 2010), which may explain the significant differences in gene expression between uninfected and infected mosquitoes at 24 hpi (Ferreira et al. 2022). In addition, Vlachou et al. (2005) using the *Plasmodium berghei* - *Anopheles gambiae* model found that 7% of the mosquito transcriptome was differentially regulated during ookinete invasion of the midgut.

At 24 hpi, for both mosquitoes infected with *P. cathemerium* and *P. relictum*, we found an increase in differentially expressed genes and enriched GO terms related with the mitochondrial respiratory chain activity that ultimately produces reactive oxygen species (ROS) (Kowaltowski et al. 2009). Small regulatory changes in the mitochondrial respiratory chain can drastically affect ROS generation (Korshunov et al. 1997; Kowaltowski et al. 2009). For example, *Anopheles stephensi* and *An. gambiae* increased the levels of ROS in response to *Plasmodium* infection (Han 2000; Kumar et al. 2003), and higher levels of ROS improve mosquito survival after a bacterial infection (Molina-Cruz et al. 2008). At 24 hpi we also found most of the differences in gene expression between mosquitoes infected by each of these parasites.

### Differential immune response between mosquitoes exposed to P. relictum or P. cathemerium

We found important differences in the expression of genes associated with the immune response between *P. relictum*- and *P. cathemerium*-infected mosquitoes. At 24 hpi, in *Cx. pipiens* infected by *P. cathemerium* compared to controls, we found that two of the most up-regulated genes were protein G12 precursors. These two genes and other three protein G12 precursor genes were down-regulated in mosquitoes infected by *P. relictum* compared to those infected by *P. cathemerium*. G12 transcripts accumulate in the midgut of mosquitoes after blood feeding (Shao et al. 2005; Bonizzoni et al. 2012), and may aid in erythrocyte digestion given their hemolytic activity (Foo et al. 2020). G12 transcripts have been suggested to play a role in immune function because: i) G12 protein in Ae*. Aegypti* has a high level of identity with cockroach allergens (Morlais et al. 2003), ii) it is up-regulated after flavivirus infection via the JAK-STAT pathway (Etebari et al. 2017), and iii) it has a cytolytic effect on flaviviruses and several types of eukaryotic cells (Foo et al. 2020). In fact, a G12 protein gene was found up-regulated in *Ae. aegypti* 12 hours after feeding on blood infected with *Plasmodium gallinaceum* (Morlais et al. 2003), suggesting a potential role in the immune response against these parasites as well.

In *P. relictum*-infected mosquitoes two of the up-regulated genes at 24 hpi were related to the Toll pathway of the innate immune response when compared to the controls: the Spätzle and Cecropin N proteins. In insects, when a pathogen is recognized, the extracellular Spätzle cytokine activates the Toll receptors, which regulate the antimicrobial peptides (AMPs), an essential innate immune response (De Gregorio 2002; Shia et al. 2009). The AMPs eventually kill pathogens by a number of strategies including disrupting the microbial membrane (Shen et al. 2018). Spätzle protein activates a Toll receptor in *Ae. aegypti* mosquitoes when infected with the fungus *Beauveria bassiana* (Shin et al. 2006) and in other insects after bacterial and fungal exposure (Bae et al. 2021). Cecropins are one of the largest groups of insect AMPs found in the orders Diptera, Lepidoptera, and Coleoptera among others (Vizioli et al. 2000; An et al. 2009; Memarpoor-Yazdi et al. 2013). The activation of the Toll pathway and specifically cecropin-analogs may kill *Plasmodium* parasites (Jaynes et al. 1988; Frolet et al. 2006) and disrupt sporogonic development by aborting the normal development of oocysts (Gwadz et al. 1989; Kim et al. 2004), which could reduce the parasite load in posterior infection stages. Our results showing the role of the Toll pathway are consistent with a recent study addressing the effect of *P. relictum* (lineage SGS1) on the immune system of *Cx. quinquefasciatus* (García-Longoria et al. 2022). They found that over 50% of immune genes identified as being part of the Toll pathway in *Cx. quinquefasciatus* were up-regulated after exposure to *P. relictum*.

### Differential expression of genes associated with trypsin and serine metabolism

Serine proteases have several functions in insects including blood digestion (Borovsky and Schlein, 1988) and mediation in the immune response (melanization, cytokine activation, and antimicrobial peptides; Jiang et al. 2010). Trypsins and chymotrypsins, two types of serine proteases, are two essential digestive enzymes in mosquitoes (Molina-Cruz et al. 2005; Borges-Veloso et al. 2012). The synthesis of trypsin and chymotrypsin is triggered after mosquito blood feeding to digest the chitin-containing peritrophic matrix (Vizioli et al. 2001; Muller et al. 1995) which is fully formed at 24 h post feeding (Hegedus et al. 2009). However, the production of trypsin and chymotrypsin may be down or up-regulated by the infection with different parasites (Borovsky and Schlein 1987; Shahabuddin et al. 1996; Serrano-Pinto et al. 2010). Mosquito trypsin may be a signal for *Plasmodium* ookinetes to cross the peritrophic matrix at the right time for proper *Plasmodium* development (Shahabuddin et al. 1996). In particular, mosquito trypsin proteases play a fundamental role in allowing *P. gallinaceum* to cross the peritrophic matrix by activating a *Plasmodium* prochitinase enzyme (Shahabuddin et al. 1996). In fact, trypsin inhibitors block the development of *Plasmodium* oocysts (Shahabuddin et al. 1996), suggesting that mosquito trypsin is a key molecule for pathogen infection.

When compared to controls, mosquitoes exposed to *P. cathemerium* up-regulated the trypsin 2 precursor gene, while mosquitoes exposed to *P. relictum* strongly down-regulated genes ofchymotrypsins and serine protease precursors . Furthermore, when comparing gene expression between mosquitoes exposed to each species, we found that mosquitoes exposed to *P. relictum* also down-regulated the chymotrypsins and serine protease precursors that when compared to control, as well as others chymotrypsins, serine protease precursors, and three trypsin precursos. This differential expression was found at 24 hpi, therefore coinciding with the moment when the peritrophic matrix is fully formed and the ookinetes are crossing it. Agreeing with the findings of Shahabuddin et al. (1996) for the infection by *P. gallinaceum*, we also found that trypsin (*trypsin 2 precursor*) was up-regulated in mosquitoes infected with *P. cathemerium* when compared with uninfected mosquitoes. This suggests the potential role of the trypsin proteases to allow *P. cathemerium* to cross the peritrophic matrix. In contrast, mosquitoes infected with *P. relictum* had a number of down-regulated genes characterized as serine proteases, including several chymotrypsin and one trypsin. Ferreira et al. (2022) also found serine-type endopeptidase activity enriched for down-regulated genes when studying the expression of *Cx. quinquefasciatus* infected by *P. relictum.* We hypothesize that the lower levels of trypsin in response to *P. relictum* infection might lead to a lower number of oocysts and would explain why the transmissibility of *P. relictum* is lower than that of *P. cathemerium* (Gutiérrez-López et al. 2020).

In addition, an under-expression of serine-proteases related to blood digestion such as chymotrypsins should partly block the digestion of the blood, which is very rich in proteins (Kumar et al. 2017). This would make it difficult to obtain nutrients needed for various processes like egg formation and could lead to digestive dysregulation in the mosquito which might increase mortality. Here mosquitoes seem to be down-regulating important processes due to the infection with *P. relictum* which might have a cost for the mosquito, induce damage and decrease the tolerance to infection.

### Reduced differential gene expression at 10 and 21 days post-infection

An unexpected result was the strong decrease of differential gene expression between infected and uninfected mosquitoes at 10 dpi. We chose this time point because at about 10 dpi *Plasmodium* mature oocysts release the sporozoites into the mosquito hemocoel (Aly et al. 2020). However, at this time point, only *P. relictum*-infected mosquitoes compared to controls showed differences in gene expression and the number of differentially expressed genes decreased significantly compared to 24 hpi. Similar results were obtained by Ferreira et al. (2022) when studying gene expression of *Cx. quinqueafasciatus* exposed to *P. relictum*. In addition, the absence of differential expression in mosquitoes exposed to *P. relictum* compared to those exposed to *P. cathemerium* may be due to differences in the developmental time of different species of *Plasmodium* (Sinden 1983). *P. cathemerium* produces sporozoites faster than *P. relictum* when infecting *Cx. pipiens* mosquitoes (Kazlauskienė et al. 2013; Aly et al. 2020), and therefore they may not be exactly at the same point of development within the mosquito at 10 dpi. In addition, when the sporozoites are realesed, part of them are destroyed by a rapid immune response (Hillyer et al. 2007), which may also result in a lower genetic response by the mosquito.

By 21 dpi, the differences in gene expression were drastically reduced. With only two genes differentially expressed in mosquitoes infected with *P. relictum*, and none in those infected with *P. cathemerium*. This decrease in differential expression towards the end of the infection has been found before in *Cx. quinqueafasciatus* infected with *P. relictum* (Ferreira et al. 2022, García-Longoria et al. 2022). Although the causes behind this decrease are not clear, we hypothesize that the response towards the end of the infection might be localized to the salivary glands, and by analysing whole mosquitoes we are diluting any potential effect.

### Comparisons across avian malaria studies

Although studies analysing the response of mosquitoes to avian malaria are still scarce we can already see some common patterns and differences. Two other recent studies have analysed the response to *P. relictum* infection in *Cx. quinquefasciatus*, a species closely related to *Cx. pipiens*. A common pattern found in all studies is the progressive reduction in the differential gene expression at 10 and 21 dpi with respect to 24 hpi. In addition, like García-Longoria et al. (2022), we found the activation of the Toll-like receptor pathway in response to *P. relictum SGS1* infection. Interestingly, Ferreira et al. (2022) did not find this pattern. Although different factors may be influencing this result, like experimental procedures or the genetic differences between the populations of *Cx*. *quinquefasciatus* used, one of the main factors might have been the lineage used for the infection. Ferreira et al. (2022) infected the mosquitoes with another lineage, *P. relictum* GRW4, which dominates in America. Like Ferreira et al. (2022), the response to infection we found for mosquitoes infected with *P. cathemerium*, was different. But, a similar pattern has been found before. Shahabuddin et al. (1996) reported the important role of a mosquito trypsin in the passage of *P. gallinaceum* through the peritrophic matrix, as our results indicate for *P. cathemerium*. Altogether these result suggests that different *Plasmodium* species or lineages may trigger a differential immune response in mosquitoes.

### Concluding remarks

Because we used naturally infected wild birds as *P. cathemerium* and *P. relictum* donors and wild collected mosquitoes, the results obtained here represent a good example of a natural system. In this respect, we found a different transcriptomic response to infections, especially at 24 hpi. This time point coincides with one of the key stages of *Plasmodium* development in mosquitos, when the ookinetes form, cross the peritrophic matrix and start to invade the midgut epithelium. Both elicit an innate immune response, with mosquitoes exposed to *P. relictum* activating the Toll Pathway (Spätzle and Cecropin N proteins) which may result in a a reduction of *Plasmodium* development (Frolet et al. 2006) and *P. cathemerium-* infected mosquitoes up-regulating the antimicrobial G12 proteins. On the other hand, the lower levels of trypsin in mosquitoes exposed to *P. relictum* compared to those expose to *P. cathemerium* may affect the parasite development within the mosquito, which may affect parasite transmission (Gutiérrez-López et al. 2020). If the cost of this response is high, this can also potentially lead to higher mosquito mortality. In particular, the proteases and trypsins are necessary to digest the blood meal, and if levels are too low this might increase mosquito mortality. Future studies are necessary to better understand how the observe differences are linked to differences in transmission. In addition, it will also be important to understand how these differences may be related to the different ecology and incidence of *Plasmodium* lineages/species in the wild.

## Supporting information

File including all supplemental figures and tables cited in the main text

## 5. Acknowledgments

We thank Francisco Ferreira for his help optimizing the RNA extraction protocol, Juan Pascual, and Alazne Diez for their help during the experiments, and Cristina Pérez for her help with the laboratory work. This publication was supported by the project Research Infrastructures for the control of vector-borne diseases (Infravec2), which has received funding from the European Union’s Horizon 2020 research and innovation program, under grant agreement No 6738; project PGC2018-095704-B-I00 from Agencia Española de Investigación supported by FEDER Funds from the European Union and the computing infrastructure provided by ICTS-RBD-CSIC. This study was also partially financed by the PID2020-118205GB-I00 grant funded by MCIN/AEI/ 10.13039/501100011033 and by “ERDF A way of making Europe”. MG was supported by a FPI grant (PRE2021-098544). GY contributions were supported by the Faculty of Biochemistry, Biophysics and Biotechnology at Jagiellonian University (Poland), under the Strategic Programme Excellence Initiative.

## 7. Data Availability Statement

Raw sequences generated in this study have been submitted to the European Nucleotide Archive ENA database (https://www.ebi.ac.uk/ena/browser/home) under project accession number PRJEB1609, Study ERP125411.

## 8. Author Contributions

María José Ruiz-López, Josué Martínez-de la Puente and Jordi Figuerola Borras designed the original study. María José Ruiz-López developed the experimental assay and the laboratory work. Marta Garrigós did the bioinformatic work and interpreted the results with María José Ruiz-López and Guillem Ylla. Marta Garrigós wrote the first original draft of the manuscript and subsequent versions with considerable assistance from the rest of the authors.

