## Supplementary material for "Two avian *Plasmodium* species trigger different transcriptional responses on their vector *Culex pipiens*": File including all supplemental figures and tables cited in the main text

**Table S1**: General information of samples and their reads, including infection status, time post-infection, number of raw reads, and number and percentage of clean reads and reads mapped to Culex quinquefasciatus. Shaded samples were removed for downstream analyses because they presented barcoding errors due to lab processing.

| Sample | Infection  status | Time | Raw reads | Clean reads (%) | Reads mapped to  *Culex quinquefasciatus* |
| --- | --- | --- | --- | --- | --- |
| T1-1-51 | Control | 24h | 33177082 | 31008365 (93,46) | 26038949  (83.98) |
| T1-2-51 | Control | 24h | 35149742 | 34069123 (96,93) | 28672552  (84.16) |
| T1-3-51 | Control | 24h | 32243129 | 30936498 (95,95) | 25993214  (84.02) |
| T1-4-51 | Control | 24h | 22466452 | 21294753 (94,78) | 16645600  (78.17) |
| T1-1-67 | *P. cathemerium* | 24h | 22498217 | 21936666 (97,50) | 18332032  (83.57) |
| T1-2-67 | *P. cathemerium* | 24h | 27397567 | 27128906 (99,02) | 22874928  (84.32) |
| T1-3-67 | *P. cathemerium* | 24h | 33324219 | 33162522 (99,51) | 27433799  (82.72) |
| T1-4-67 | *P. cathemerium* | 24h | 26626804 | 26462620 (99,38) | 21953450  (82.96) |
| T1-1-85 | *P. relictum* | 24h | 30674241 | 30435998 (99,22) | 24801694  (81.49) |
| T1-2-85 | *P. relictum* | 24h | 23278856 | 22883101 (98,30) | 19001869  (83.04) |
| T1-3-85 | *P. relictum* | 24h | 35771588 | 34263353 (95,78) | 28608913  (83.50) |
| T1-4-85 | *P. relictum* | 24h | 30472866 | 29351327 (96,32) | 24186857  (82.41) |
| T2-5-51 | Control | 10d | 28177843 | 25003834 (88,74) | 20239066  (80.95) |
| T2-16-51 | Control | 10d | 20967331 | 18808738 (89,70) | 15330448  (81.51) |
| T2-21-51 | Control | 10d | 25806437 | 22496937 (87,18) | 18230162  (81.04) |
| T2-15-51 | Control | 10d | 15406828 | 12411723 (80,56) | 9924380  (79.96) |
| T2-13-67 | *P. cathemerium* | 10d | 27298353 | 25826907 (94,61) | 19709378  (76.31) |
| T2-16-67 | *P. cathemerium* | 10d | 29882582 | 28545072 (95,52) | 21510841  (75.36) |
| T2-23-67 | *P. cathemerium* | 10d | 25577894 | 25107302 (98,16) | 19754366  (78.68) |
| T2-24-67 | *P. cathemerium* | 10d | 29176625 | 29037230 (99,52) | 22315395  (76.85) |
| T2-13-85 | *P. relictum* | 10d | 37005797 | 35991229 (97,26) | 29906455  (83.10) |
| T2-16-85 | *P. relictum* | 10d | 29891263 | 28936485 (96,81) | 24051388  (83.12) |
| T2-5-85 | *P. relictum* | 10d | 34277655 | 32255293 (94,10) | 25436232  (78.86) |
| T2-6-85 | *P. relictum* | 10d | 32903444 | 31841055 (96,77) | 26529969  (83.32) |
| T3-11-51 | Control | 21d | 32385022 | 32118665 (99,18) | 26043488  (81.09) |
| T3-17-51 | Control | 21d | 32770699 | 32039074 (97,77) | 26111299  (81.50) |
| T3-18-51 | Control | 21d | 29728686 | 29263504 (98,44) | 20838799  (71.21) |
| T3-21-51 | Control | 21d | 30766572 | 30159889 (98,03) | 24335913  80.69) |
| T3-17-67 | *P. cathemerium* | 21d | 21530168 | 21145476 (98,21) | 16532428  (78.19) |
| T3-18-67 | *P. cathemerium* | 21d | 37716163 | 37090714 (98,34) | 30680752  (82.72) |
| T3-22-67 | *P. cathemerium* | 21d | 28765518 | 28497563 (99,07) | 23298862  (81.76) |
| T3-9-67 | *P. cathemerium* | 21d | 39203774 | 35121618 (89,59) | 27087303  (77.12) |
| T3-10-85 | *P. relictum* | 21d | 45332626 | 44887231 (99,02) | 36489870  (81.30) |
| T3-11-85 | *P. relictum* | 21d | 33120712 | 32798179 (99,03) | 27092510  (81.90) |
| T3-14-85 | *P. relictum* | 21d | 41161560 | 40739559 (98,97) | 32717880  (80.31) |
| T3-9-85 | *P. relictum* | 21d | 28181782 | 28034675 (99,48) | 21149517  (80.18) |

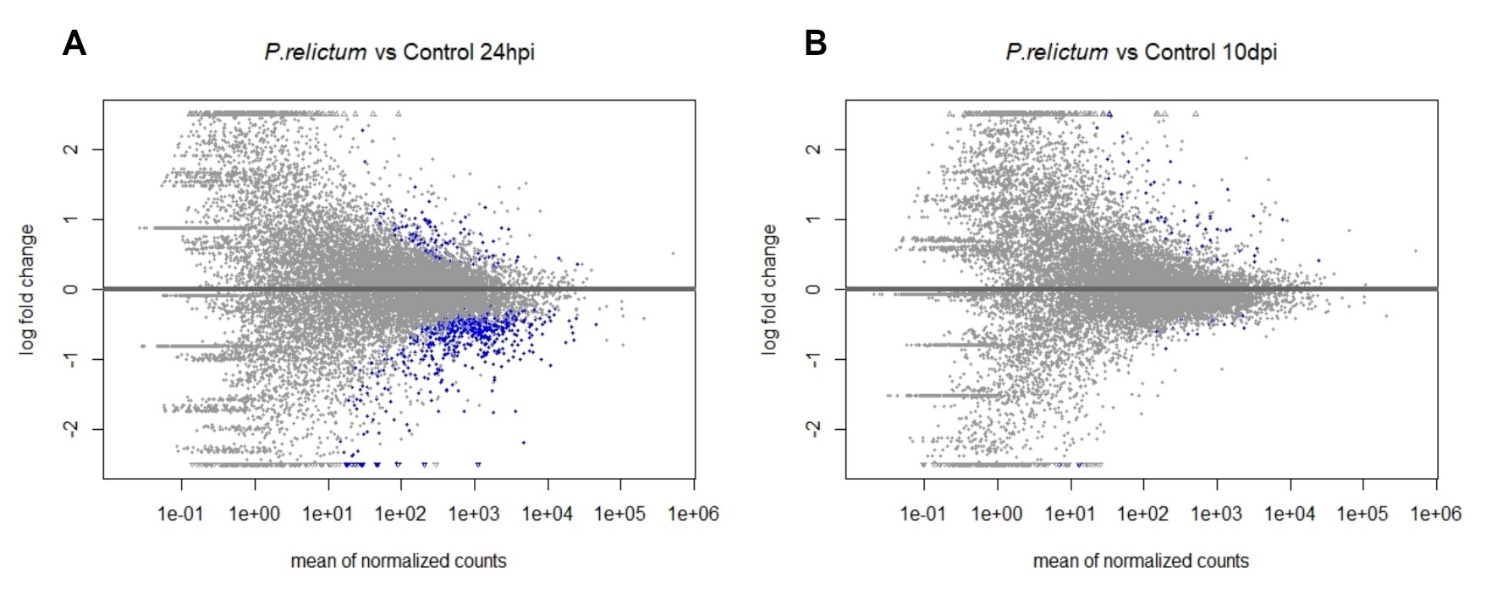

**Figure S1.** MA plot for Plasmodium relictum vs controls gene expression at (A) 24 hours post infection and (B) 10 days post infection. Triangles correspond to genes with a log fold change greater than 1 or less than -1. Blue dots and triangles correspond to genes considered differentially expressed (adjusted p-value < 0.01). The y-axis shows the log fold change (i.e. the intensity of the gene expression). The y-axis shows the average expression between the groups compared.

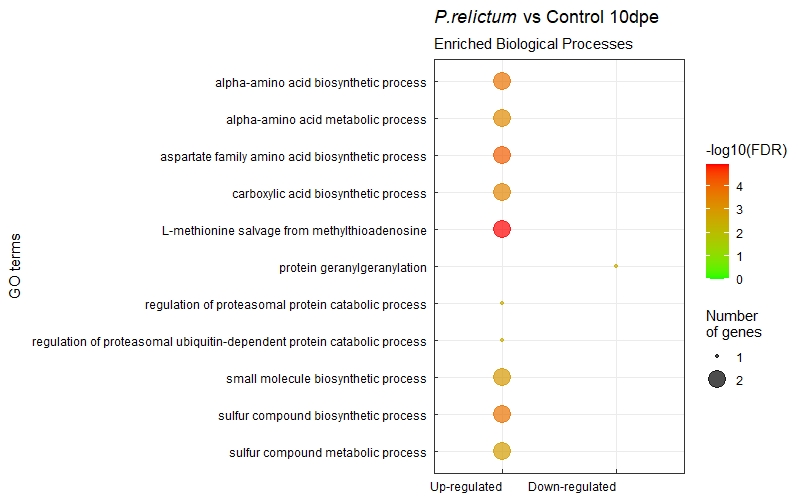

**Figure S2.** Dot plot of enriched GO biological processes for differentially expressed genes (up and down-regulated) in P. relictum infected mosquitoes vs controls at 10 dpi. The y-axis lists the GO terms ordered by Fisher exact p-value. The legend shows both the number of significant genes (dot size) and the –log(p-value) (color gradient). Larger dots correspond to a higher number of significant genes, and the color gradient goes from green for the least significant terms to red for the most significant terms.

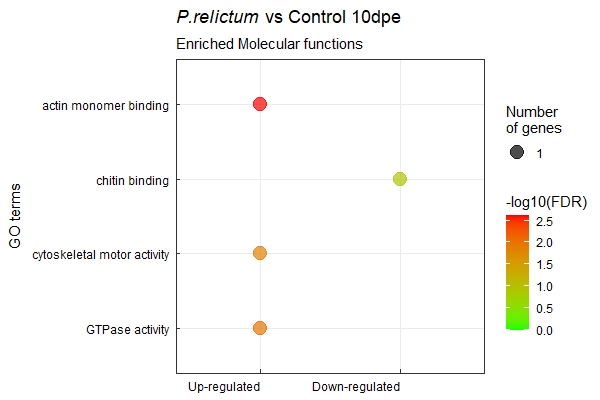

**Figure S3**. Dot plot of enriched GO molecular functions for differentially expressed genes (up and down-regulated) in P. relictum infected mosquitoes vs controls at 10 dpi.

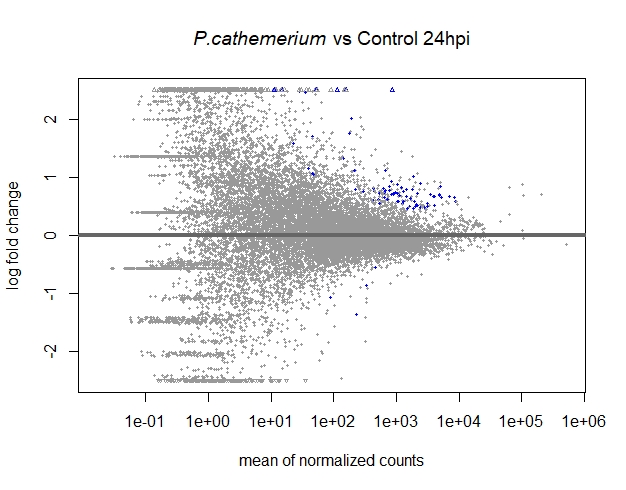

**Figure S4**. MA plot for *P. cathemerium* vs controls gene expression at 24 hpi.

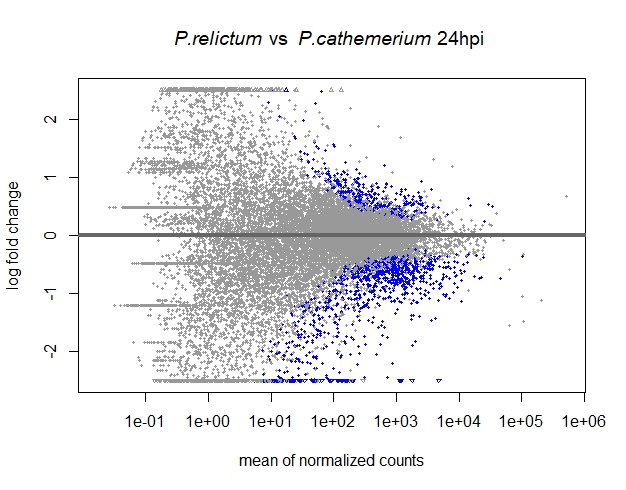

**Figure S5**. MA plot for P. relictum vs P. cathemerium gene expression at 24 hpi.
